# Small circRNAs with self-cleaving ribozymes are frequently expressed in metazoan transcriptomes

**DOI:** 10.1101/721605

**Authors:** Amelia Cervera, Marcos de la Peña

**Affiliations:** IBMCP (CSIC-UPV). C/ Ingeniero Fausto Elio s/n, 46022, Valencia, Spain

**Keywords:** circular RNA, retrotransposons

## Abstract

Ribozymes are catalytic RNAs present in modern genomes but considered as remnants of a prebiotic RNA world. The paradigmatic hammerhead ribozyme (HHR) is a small self-cleaving motif widespread from bacterial to human genomes. Here, we report that most of the classical type I HHRs frequently found in the genomes of diverse animals are contained within a novel family of non-autonomous non-LTR retrotransposons. These retroelements are expressed as abundant linear and circular RNAs of ∼170-400 nt in different animal tissues. *In vitro* analyses confirm an efficient self-cleavage of the HHRs harboured in invertebrate retrozymes, whereas those in retrozymes of vertebrates, such as the axolotl, require to act as dimeric motifs to reach higher self-cleavage rates. Ligation assays of retrozyme RNAs with a protein ligase versus HHR self-ligation indicate that, most likely, tRNA ligases and not the ribozymes are involved in the step of RNA circularization. Altogether, these results confirm the existence of a new and conserved pathway in animals and, likely, in eukaryotes in general, for the efficient biosynthesis of RNA circles through small ribozymes, which will allow the development of biotechnological tools in the emerging field of circRNAs.

## INTRODUCTION

The discovery of catalytic RNAs or ribozymes in the 80’s (1,2) strongly supported the hypothesis of a prebiotic RNA world, where the first living organisms were based on RNA as both the genetic material and as catalyst (3-5). Some of these ancient ribozymes are thought to subsist in modern organisms, carrying out crucial roles such as peptide bond formation (6), tRNA processing (2) or mRNA splicing (7). There is, however, an enigmatic group of small (∼50-150 nt) self-cleaving ribozymes with uncertain origins and functions (8,9). The hammerhead ribozyme (HHR), the first and best-studied member of this family, has a conserved core of 15 nucleotides surrounded by three helixes (helix I to III), which adopt a γ-shaped fold where helix I interacts with helix II through characteristic tertiary interactions required for efficient self-cleavage (10-12). There are three possible circularly permuted forms named type I, II or III depending on the open-ended helix of the motif (Figure 1A). The HHR was firstly discovered in infectious circRNAs of plants, such as viral RNA satellites and viroids (13), but also in the repetitive DNA of some amphibians (14), and, later on, in schistosomes (15) and cave crickets (16). A few years ago, however, HHR motifs were found ubiquitously from bacterial to eukaryotic genomes (17-21), including humans (22), unveiling this catalytic RNA as the most widespread small ribozyme. The biological roles of these genomic HHRs, however, have remained poorly known. Rare minimal variants of the type I HHR (Figure 1A) have been reported in retrotransposons of the Penelope-like (PLEs) (23) and Terminon (24) families, whereas a few copies of highly conserved HHRs in amniotes seem to play a role in mRNA biogenesis (25). More recently, type III HHRs detected in several flowering plants have been found to be involved in the processing of a novel family of non-autonomous LTR retrotransposons, the so-called *retrozymes* for *retro*element with hammerhead ribo*zymes* (Figure 1B), which spread through circRNA transposition intermediates of 600-800 nt (26).

In this work, the classical type I HHR motifs reported since 1987 in diverse metazoan genomes have been thoroughly studied. Analyses were performed with tissues of animals from three distant phyla: cnidaria (a coral), mollusca (a mussel) and chordata (a salamander). Our results are consistent with a conserved role for most type I HHRs in the life cycle of a novel family of constitutively expressed non-autonomous non-LTR retrolements, which spread thoughout animal genomes by means of small circRNAs transposition intermediates.

**Figure 1.**
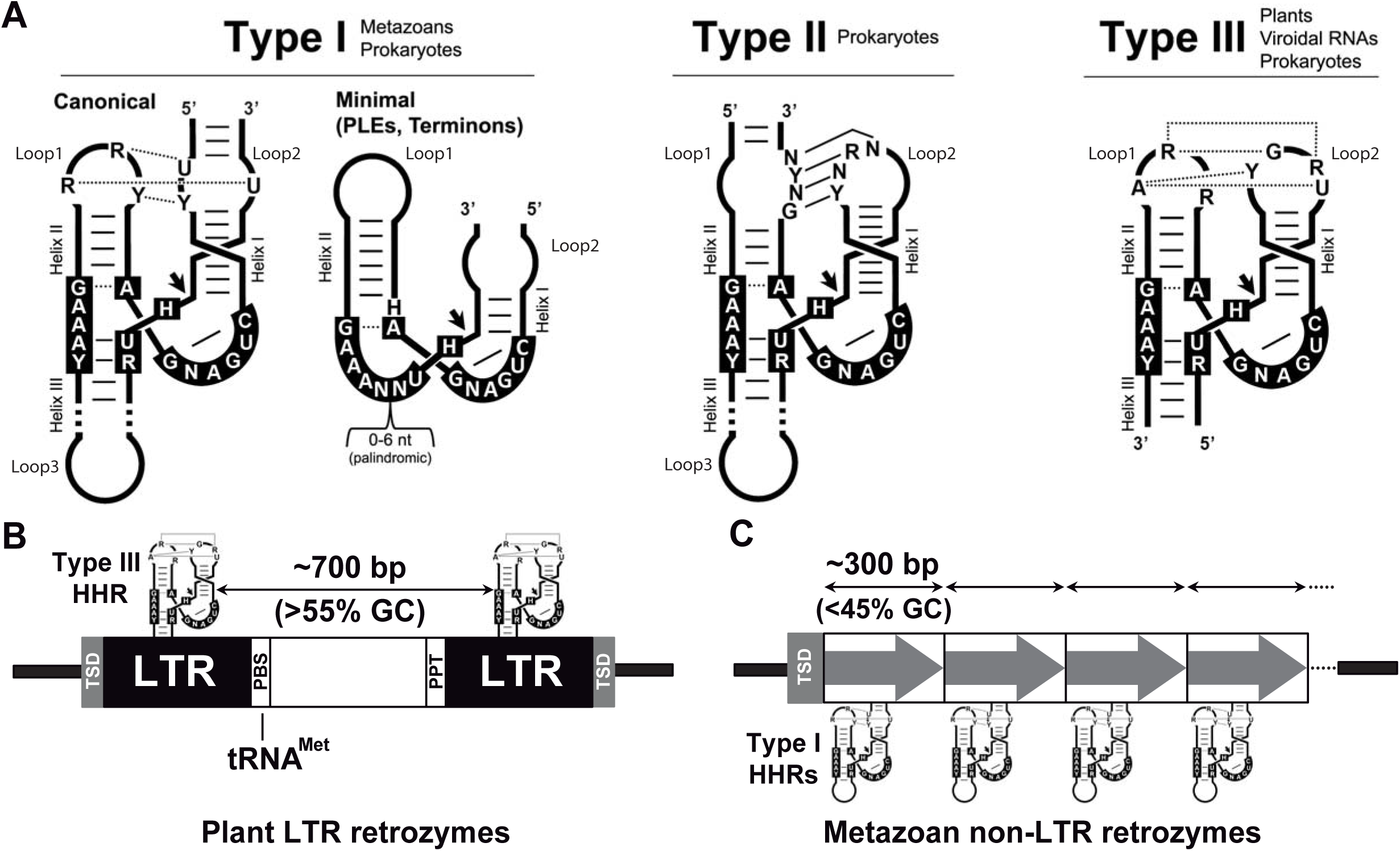
(A) Schematic representation of the three possible hammerhead ribozyme (HHR) topologies (Type I, II and III). The most frequent nucleotides in the catalytic core (black boxes) and in the loop-loop interactions are indicated. Dotted and continuous lines refer to Non-canonical and Watson-Crick base pairs. Black arrows indicate the self-cleavage site. The three HHR types have been reported in prokaryotic/phages genomes, whereas metazoan and plant genomes mostly show type I and III motifs, respectively. (B) Schematic representation of a typical LTR retrozyme from plant genomes containing type III HHRs. The average size encompassed by the HHRs (∼700 bp) and GC content (>55%) are indicated (C) Canonical type I HHRs in metazoan genomes can be found within short tandem repeats (∼300 bp, from dimers to large multimers) of low GC content (<45%), which can be regarded as a new family of non-LTR retrozymes.

## MATERIALS AND METHODS

### Bioinformatics

The RNAmotif software (27) was used for the detection of both canonical and minimal HHR motifs (either type-I or type-III architectures) in metazoan genomes (RefSeq Genome Database, invertebrate/ and vertebrate_other/, plus some other individual genomes based on the obtained results). The hits obtained were inspected for the presence of tertiary interactions between helix I and II to ensure they were *bona fide* HHRs, and their occurrence within tandem repeats. Sequence homology searches through BLAT (28). BLAST and BLASTX (29) tools were carried out in individual or grouped genomes. Sequence alignments were performed with ClustalX (30) and Jalview (31) software. Secondary RNA structures of minimum free energy were calculated with the RNAfold program from the ViennaRNA Package (32) and depicted with RNAviz (33).

### Nucleic acid extractions

DNA from animal tissues was extracted following a CTAB-based protocol with some modifications (26). Briefly, samples were homogenised (1:2.5 w/v) in extraction buffer (1% w/v CTAB, 50 mM Tris-HCl pH 8, 10 mM EDTA, 0.7 M NaCl, 1% w/v PVP-40, 1% 2-mercaptoethanol), and were incubated for 30 min at 60°C. An equal volume of chloroform:isoamyl alcohol (24:1 v/v) was added to the samples, which were then centrifuged. DNA in the supernatant was precipitated with 0.1 volumes of 3 M sodium acetate (pH 5.5) and 2.5 volumes of 100% ethanol, dissolved in MilliQ water (Millipore), and quantified in a NanoDrop 1000 Spectrophotometer (Thermo Fisher Scientific).

RNA from animal tissues was isolated with Trizol reagent (Invitrogen), following the manufacturer’s instructions. Shortly, tissues were homogenised (1:10 v/v) in Trizol, and then 0.2 volumes of chloroform:isoamyl alchohol (24:1 v/v) were added to the samples. The mixture was centrifuged, and RNA in the supernatant was precipitated with 0.1 volumes of 3 M sodium acetate (pH 5.5) and 2.5 volumes of 100% ethanol. RNA was resuspended and quantified as explained above.

### PCR, RT-PCR and molecular cloning

Genomic retrozymes were amplified by PCR using degenerate primers designed to bind the most conserved motifs within a species. The hot-start, proof-reading PrimeStar HS DNA Polymerase (Takara) was used following the manufacturer’s instructions, together with phosphorylated (T4 polynucleotide kinase, Takara) primers that spanned the length of the RNA of the retrozyme (i.e., the retrozyme monomer). Amplification products of the adequate size were extracted from native 5% 1× TAE polyacrylamide gel slices with phenol:chloroform:isoamyl alcohol (25:24:1), and concentrated by ethanol precipitation as described above.

For circular retrozyme RNA cloning by RT-PCR, RNA was fractionated by native 5% 1× TAE PAGE, and extracted from gel slices corresponding to the retrozyme size (300 – 500 nt for *M. galloprovincialis*). RNAs were reverse-transcribed with SuperScript II (Invitrogen), and PCR-amplified with adjacent divergent primers previously phosphorylated, using Prime Star HS DNA polymerase.

PCR products were cloned into a linearised pBlueScript KS (Agilent) vector by blunt-end ligation, and were sequenced automatically with an ABI Prism DNA sequencer (Perkin–Elmer).

### RNA transcriptions

RNAs of the cloned retrozymes were synthesized by *in vitro* run-off transcription of pBlueScript KS plasmids containing the corresponding retrozyme fragment or HHR motif linearized with the appropriate restriction enzyme. Transcription reactions contained: 40 mM Tris–HCl, pH 8, 6 mM MgCl2, 2 mM spermidine, 0.5 mg/ml RNase-free bovine serum albumin, 0.1% Triton X-100, 10 mM dithiothreitol, 1 mM each of CTP, GTP and UTP, 0.1 mM ATP plus 0.5 µCi/µl [α-32P]ATP, 2 U/µl of human placental ribonuclease inhibitor, 20 ng/µl of plasmid DNA, and 4 U/µl of T7 or T3 RNA polymerases. After incubation at 37 °C for 2 h, the products were fractionated by polyacrylamide gel electrophoresis (PAGE) in 5% (retrozymes) or 15% gels (ribozymes) with 8 M urea.

### Kinetic analysis of self-cleavage under co-transcriptional conditions

Analyses of retrozyme self-cleavage under co-transcriptional conditions were performed as previously described (34). Transcription reactions were carried out at 37°C as described before, and appropriate aliquots (smaller volumes were taken at longer incubation times) were removed at different time intervals, quenched with a fivefold excess of stop solution at 0°C, and analysed as previously described (34). Briefly, the uncleaved and cleaved transcripts were separated by PAGE in 5% denaturing gels. The product fraction at different times, Ft, was determined by quantitative scanning of the corresponding gel bands and fitted to the equation F_t_ = F_∞_(1 - e^−kt^), where F_∞_ is the product fraction at the reaction endpoint, and k is the first-order rate constant of cleavage (k_obs_).

### Northern blot analysis

For northern blot analysis, 5 to 50 μg of purified RNA from the different animal tissues were examined in 5 % polyacrylamide gels under native (1×TAE) or denaturing (8 M urea, at either 0.25×TBE or 1×TBE) conditions. After ethidium bromide staining, RNAs were electroblotted onto nylon membranes (Amersham Hybond-N, GE Healthcare) and UV-fixed with a crosslinker (UVC 500, Hoefer). Prehybridization, hybridization (at 68°C in 50% formamide) and washing was done following the instructions of the manufacturer (GE Healthcare). The riboprobes were obtained by rub-off transcriptions of linearized pBlueScript plasmids containing the full retrozymes in the presence of DIG-UTP (Roche Diagnostic GmbH) (26).

### RNA circularization experiments

For *in vitro* analysis of the tRNA ligation and self-ligation capabilities of RNA retrozymes, monomeric retrozyme RNAs that resulted from double self-cleavage after transcription (either in the presence or in the absence of [α-32P]ATP) of dimeric constructs were purified from 5 % polyacrylamide gels under denaturing conditions (8 M urea, 1×TBE). For the assays of RtcB ligation, 1 to 10 ng of gel-purified retrozyme RNA were ligated using RtcB ligase in its corresponding buffer for 1 h at 37 °C (New England Biolabs). Best results for self-ligation assays were obtained with 1 to 10 ng of gel-purified retrozyme RNA in 50 mM Tris–HCl, pH 8, which were firstly denatured at 95 °C for 1 min, slowly cooled down to 25 °C (1 °C decrease every 2 seconds), and then incubated in the presence of 10 to 50 mM MgCl_2_ for 1h at 25 °C.

## RESULTS

### Type I HHRs in cnidarian genomes occur within small DNA tandem repeats, which are expressed as linear and circular RNAs

Previous studies have reported that many type I HHRs in animal genomes can be found within short tandem repeats of a few hundred base pairs (150 to 450 bp), widely but patchily distributed among most metazoan phyla (25). In contrast to plant LTR retrozymes, which harbour type III HHRs in a dimeric arrangement (Figure 1B), metazoan repeats with type I HHRs occur in multiple copies and lack the characteristic long terminal repeats (LTRs), primer binding site (PBS), and polypurine tract (PPT) motifs. These sequence repeats in animals show a low GC content (below 50%), and, when present, target side duplications (TSDs) are larger than the typical 4 bp TSDs found in plant LTR retrozymes (26) (Figure 1B). Altogether, these data suggest that metazoan repeats with type I HHRs constitute a family of non-autonomous non-LTR retrozymes comparable to the family of plant LTR retrozymes (Figure 1C).

To get a deeper insight of the biology of these putative retroelements with type I HHRs, we initially performed a bioinformatic analysis of the genome of the coral *Acropora millepora* (35), as an example of a low-complexity animal. This stony coral shows the presence of more than 6,000 type I HHR motifs (either minimal or canonical), which, based on their primary sequence can be classified into 16 different families. Up to 1,508 of these motifs correspond to canonical type I HHRs, showing typical helix sizes and tertiary interactions between loops 1 and 2 (12), whereas all the other motifs resembled to the atypical minimal HHR variants found in PLEs and Terminon retrotranposons (23,24) (supplementary Figure S1). The canonical *A. millepora* HHRs can be detected within small (from 220 to 240 bp) tandem repeats of the kind mentioned above, ranging from dimers and trimers, up to 62-mers. A genomic copy of a monomeric retrozyme was amplified with adjacent oligos, cloned and transcribed *in vitro.* The HHR contained in the genomic retrozyme showed an efficient RNA self-cleavage, with a co-transcritpional k_obs_ of ∼0.8 min^-1^ (Figure 2A). Northern blot analysis of *A. millepora* RNA extracts (∼30μg) from different coral samples run in polyacrylamide gels (PAGE) under native conditions showed the presence of a single band above 200 nt size, which corresponds to the size of the repeats found in the genome (Figure 2B, left). Based on the hybridization signal of a retrozyme transcript marker (0.1 ng) included in the gels, it can be estimated that the detected retrozyme RNA in coral extracts represents ∼0.1‰ of the total RNA in the sample. When the same extracts were run under denaturing conditions, the single retrozyme RNA bands observed under native conditions split into linear (∼220-240 nt) and slower migrating bands (∼300 nt), a characteristic behaviour of circular RNA molecules run in PAGEs under denaturing conditions (Figure 2B, centre). An increased and clearer separation of the linear and circular (∼400 nt size) forms of the retrozyme RNA was obtained when the ionic strength of the gel was lowered under equivalent denaturing conditions (Figure 2B, right) (36). The circular nature of part of the population of retrozyme RNAs was confirmed by RT-PCR amplification of PAGE-purified RNAs. These retrozyme circRNAs are predicted to adopt a stable and highly self-paired secondary structure (Figure 2C). Two major sequence variants of the *A. millepora* retrozyme monomer can be detected in the coral genome, with around 225 and 245 bp due to a typical insertion of 20 nucleotides, among other smaller indels along the sequence of the repeats. Previous (26) and new sequence analyses of the genomes of other stony corals show similar retrozyme sequences (supplementary Figure S2), which indicates a phylogenetic relationship among all of them. Moreover, small repeats of ∼230-350 bp carrying type I HHRs can be also identified in the genomes of diverse species of jellyfish (Figure 2D) and sea anemones (supplementary Figure S3), indicating that small circRNAs with HHRs are frequently encoded in the genomes of cnidarians.

**Figure 2.**
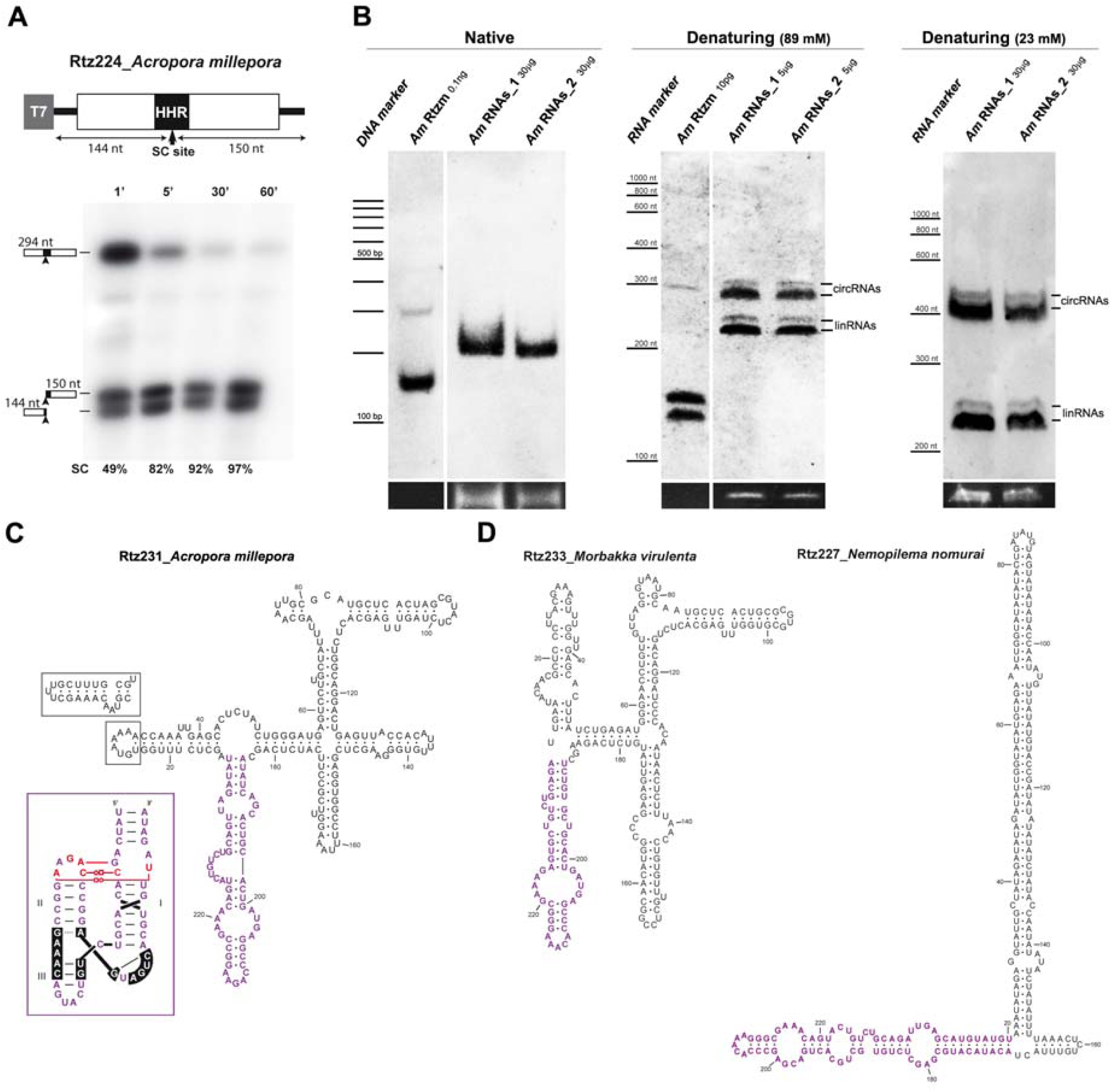
(A) A genomic retrozyme copy from the coral *A. millepora* was cloned 3’ to a T7 promoter. Schematic representation of the clone and the expected sizes of the fragments after HHR self-cleavage are indicated. Below, autoradiography of a run-off transcription of this construct at different times run in a denaturing gel. The volumes of the samples were adjusted to show an equivalent signal. RNA self-cleavage of the retrozyme reaches almost completion after 5 minutes. (B) Northern blot analysis of RNA extracts from two different samples of the coral *Acropora millepora* (Am_RNAs_1 and Am_RNAs_2) and a quantified RNA from the *in vitro* transcribed *A. millepora* retrozyme (Am Rtzm) under native (left membrane) and denaturing conditions (centre and right membranes, at 89 mM and 23 mM TBE, respectively). The approximate positions of the bands corresponding to the DNA 100-1,000bp (native blots) and RNA LR RiboRuler (denaturing blots) markers, and the linear and circular RNAs are indicated. Ethidium bromide staining of the 5S rRNAs are shown at the bottom as loading controls. (C) A minimum free energy secondary structure prediction of the circRNA derived from a representative *A. millepora* retrozyme. A typical 20 nt insertion found in many genomic retrozymes is indicated. The sequence of the HHR motif is shown in purple letters. The expected 3D structure of the HHR is shown in an inset. Non-canonical tertiary interactions between loops 1 and 2 are shown in red (50). (D) Minimum free energy secondary structure prediction for some of the retrozyme sequences detected in the genomes of the jellyfish *Morbakka virulenta* (left, 233 nt) and *Nemopilema nomurai* (right, 227 nt).

### Retrozymes with type I HHRs are expressed as heterogeneous circRNAs in mussels and, likely, many other invertebrates

Type I HHRs within short tandem repeats are also detected in the genomes of more complex metazoans, from molluscs to arthropods. The genome of the Mediterranean mussel *Mytilus galloprovincialis* (37) contains around 5,000 copies of a type I HHR, which were classified into two ribozyme families of similar sequence, but without showing most of the conserved nucleotides involved in the archetypical tertiary interactions between loops 1 and 2 (12) (Figure 1A and supplementary Figure S4). These HHRs were usually located within sequence repeats of 350-390 bp. A genomic repeat was cloned, and co-transcriptional assays revealed slower self-cleavage rates than the ones observed for the HHR in the coral retrozyme (k_obs_ ∼ 0.1 min^-1^. Figure 3A). Northern blot analyses of different mussel tissues confirmed the presence of high levels of circRNAs (up to 1‰ of the total mussel RNA) derived from these retrozyme repeats, most notably in gonads and mantle (Figure 3B). The levels of the corresponding linear RNAs of the mussel retrozymes were much lower (below 5% of the detected RNA) than the ones observed in extracts of the coral *A. millepora* (around 50%. Figure 2). However, northern assays performed with new extracts from other mussel specimens revealed higher ratios of lin/circ RNAs (up to 10-25%. Figure 3C), suggesting that the linear retrozyme RNAs detected in our assays may depend not only on the analysed tissue and species, but also on the experimental procedure during RNA extraction and manipulation.

**Figure 3.**
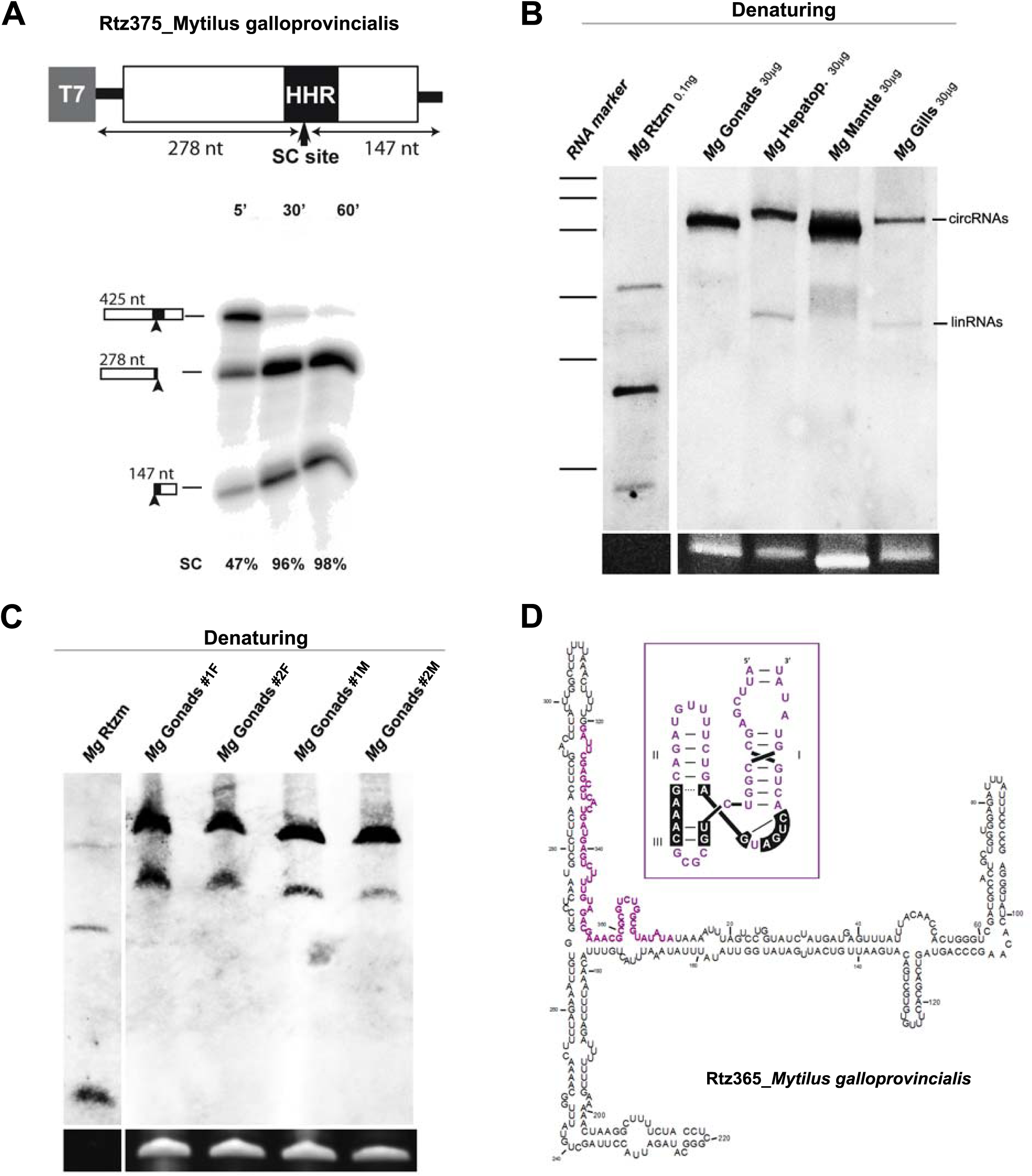
(A) Schematic representation of a genomic copy of the mussel *M. galloprovincialis* retrozyme cloned after a T7 promoter. The expected sizes of the fragments after HHR self-cleavage are indicated. Below, and autoradiography of a run-off transcription of this construct at different times run in a denaturing gel. The volumes of the samples were adjusted to show an equivalent signal. (B) Northern blot analysis under denaturing conditions of RNA extracts from different tissues of the mussel *Mytilus galloprovincialis* (gonads, hepatopancreas, mantle and gills). Mostly circular RNAs are detected. (C) Northern blot analysis under denaturing conditions of RNA extracts from gonads of different *M. galloprovincialis* mussels (two females, #1F and #2F, and two males, #1M and #2M) showing circular and, to a lesser extent, linear retrozyme RNAs. Quantified amounts of retrozyme transcripts (Mg Rtzm), size markers and loading controls are shown as in Figure 2. (D) Minimum free energy secondary structure predicted for a cloned retrozyme circRNA from mussel gonads, and its corresponding HHR structure (inset in purple)

RT-PCR and cloning of purified circRNAs from the gonads of two different mussel specimens revealed a quite heterogeneous population of retrozyme RNA sequences and sizes (supplementary Figure S5). Each different circRNA could likely result from the expression of a slightly different genomic repeat. None of the cloned retrozyme sequences, however, exactly matches any of *M. galloprovincialis* database entries, which suggests that, as previously reported for plant LTR retrozymes (26), other explanations to this sequence heterogeneity, such as RNA editing or RNA-RNA replication among others, could be considered.

Previous (25,26) and present bioinformatic analyses confirm the presence of analogous small repeats with type I HHRs, retrozyme-like, in many other invertebrate genomes from different phyla, such as rotifers, platyhelminths, annelids, crustaceans or insects, among others. Whereas the precise sequences and sizes of these tandem repeats, including the type I HHR motif, are species-specific, they all share the main features described above for non-LTR retrozymes (Figure 1C). Altogether, our data strongly suggest that metazoan retrozymes are intensively transcribed in most invertebrate tissues as small (∼170-400 nt) and highly self-paired circRNAs (supplementary Figure S6). Moreover, retrozyme containing species such as mussels and other invertebrates also show the presence of autonomous PLEs retroelements carrying atypical type I HHR variants (23,37), supporting the genomic co-existence of both families of non-autonomous and autonomous retroelements with HHRs.

### The genome of the Mexican axolotl contains up to 120,000 copies of retrozymes from two different families

The first type I HHR reported in a metazoan genome was described in 1987 in the so-called *satellite DNA* repeats (∼330 bp) of newts (14). Further studies confirmed the presence of similar transcriptionally active repeats in the genomes of different species of newts and salamanders (38). Molecular characterization of those transcription products revealed the presence of monomeric and multimeric RNAs, which were considered as linear molecules carrying 5’-hydroxyl and phosphate blocked 3’ ends (38,39). Due to a short helix III, amphibian HHRs show a low catalytic activity as single motifs, but they were found to efficiently self-cleave as dimers (40). Our bioinformatic analyses of the recently sequenced genome of the Mexican axolotl (*Ambystoma mexicanum*) (41) showed the presence of 126,247 motifs of type I HHRs similar to those previously reported in other salamander species. The axolotl ribozymes can be classified into two different groups, which occur in a comparable proportion along the genome (HHR_Amex1 and HHR_Amex2, ∼60,000 motifs each one, Figure 4A). The HHR_Amex1 motif highly resembles the type I HHRs reported in the satellite DNA of amphibians. These type I HHRs show a very weak and characteristic helix III, with a single base-paired stem capped by a palindromic tetraloop, and all the conserved tertiary interactions between loops 1 and 2. The HHR_Amex2 motifs, however, show a more stable helix III (stem with two base pairs capped by a tetraloop, similar to invertebrate HHRs), but a weaker helix II (a U U mismatch at the base of the stem), and a likely less stable tertiary interaction between loops 1 and 2 (Figure 4A). The HHR_Amex1 and HHR_Amex2 motifs are each one associated with a slightly different tandem repeat of 330 and 350 bp, respectively (supplementary Figure S7). A genomic repeat of each class (Rtz_331 and Rtz_353) was cloned and transcribed *in vitro*. As expected, transcriptional assays with the Rtz_331 monomer, harbouring a single copy of the HHR_Amex1 ribozyme, show a poor self-cleaving activity (6% self-cleavage after one hour) (Figure 4B). On the other hand, Rtz_353 carrying the HHR_Amex2 has a slightly higher self-cleaving activity during transcritpion (22% self-cleavage after one hour), although still far from the levels of self-cleavage observed for HHRs in invertebrate retrozymes (Figure 2A, 3A and 4B). Whereas the axolotl ribozymes show weak self-processing activity within monomeric repeats (i. e., with HHR acting as a monomer), dimeric constructs, where HHRs could act as dimers, result in much more efficient self-cleavage (Figure 4C), indicating that both HHR motifs may work *in vivo* as dimers. As previously described for other salamanders (39), northern blots of axolotl RNAs carried out under native conditions confirm the presence in all analysed tissues of monomeric, but also multimeric RNA repeats (Figure 4D, left). When the same RNA extracts were analysed under denaturing conditions, however, we confirmed that the monomeric RNAs that run as a single band in native conditions, are composed of a mixture of both linear and circular RNA molecules corresponding to around 330 and 350 nt (Figure 4D, right). Circular RNAs and their linear counterparts in all three analysed tissues seem to occur in a similar ratio (∼50% each). However, different accumulation levels of each variant can be readily appreciated for each tissue (Figure 4D), indicating than both 330 and 350 express differentially depending on the animal tissue. Finally, as observed for invertebrate genomes with non-LTR retrozymes (see below), the axolotl also contains numerous PLE retroelements (41) carrying minimal HHR copies in tandem (supplementary Figure 7B).

**Figure 4.**
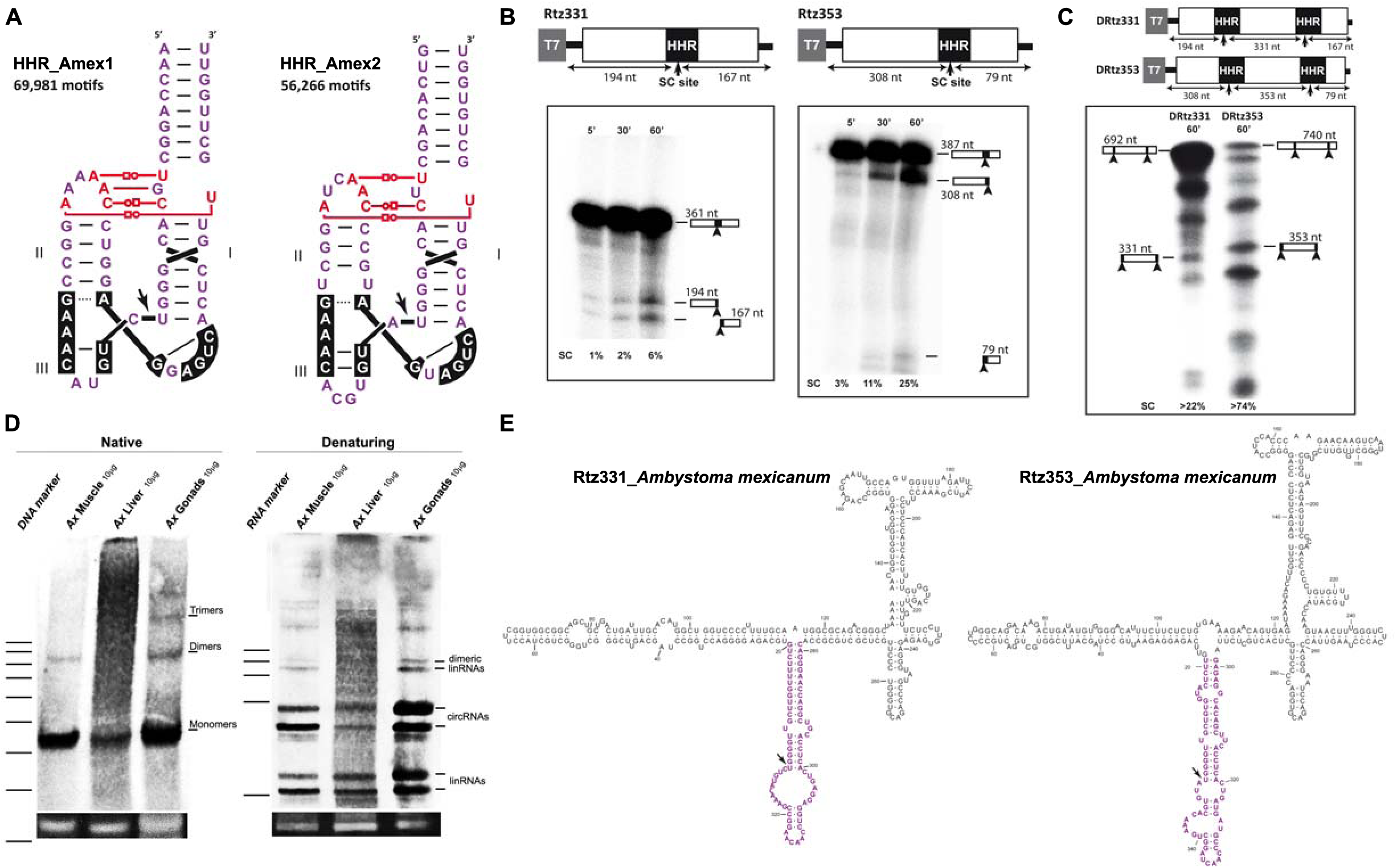
(A) The two families of canonical type I HHRs found in the genome of the axolotl (HHR_Amex1, left, and HHR_Amex2, right). The number of copies found for each motif is indicated. (B) Autoradiographies of run-off transcriptions from two genomic retrozyme copies (Rtz331 carrying HHR_Amex1 at the left, and Rtz353 carrying HHR_Amex2 at the right) of the axolotl at different times. (C) Autoradiography of 1h run-off transcriptions of the corresponding dimeric constructs shown in panel B (DRtz331 and DRtz353). (D) Northern blot analyses of RNA extracts from different axolotl tissues (muscle, liver and male gonads) that were run under native (left membrane) and denaturing conditions (right membrane). Monomeric, multimeric and circular RNAs are indicated. Quantified amounts of retrozyme transcripts, size markers and loading controls are shown as in Figure 2. (E) Minimum free energy secondary structure predictions for axolotl circRNAs of 331 and 353 nt derived from genomic retrozymes.

### *In vitro* circularization of RNA retrozymes

Natural type I HHRs keeping tertiary interactions between loops 1 and 2 have been previously reported to achieve a 2000-fold increase in the rate of ligation compared to a minimal hammerhead without the loop-loop interaction, suggesting that HHR may be almost as efficient at ligation as it is at cleavage. We previously described that RNAs encoded by plant LTR retrozymes with type III HHRs are efficiently circularized *in vitro* by a chloroplast tRNA ligase (26), similarly to the RNA circularization reported for HHR viroids (42). To get a deeper insight into the circularization mechanism of the RNAs expressed by metazoan retrozymes, linear monomers obtained *in vitro* from the *A. mexicanum* DRtzm_350 construct (Figure 4C) were assayed for either self- or tRNA ligase-mediated circularization. Although self-circularization can be readily detected, the final levels of circRNA are very low (3% of total RNA) compared with the ones obtained after incubation with the RtcB tRNA ligase (>80%) (Figure 5). Similar results were obtained with a full monomer of *M. galloprovincialis* obtained from a dimeric construct of Rtz375_Mgal, which show low levels of self-ligation (∼1%) compared with its circularization through RtcB tRNA ligase (∼30%) (Supplementary Figure S8). These observations indicate that, most likely, a tRNA ligase is also responsible for retrozyme RNA circularization in animals, in spite of the known capabilities of the type I HHRs for *in vitro* self-ligation (43).

**Figure 5.**
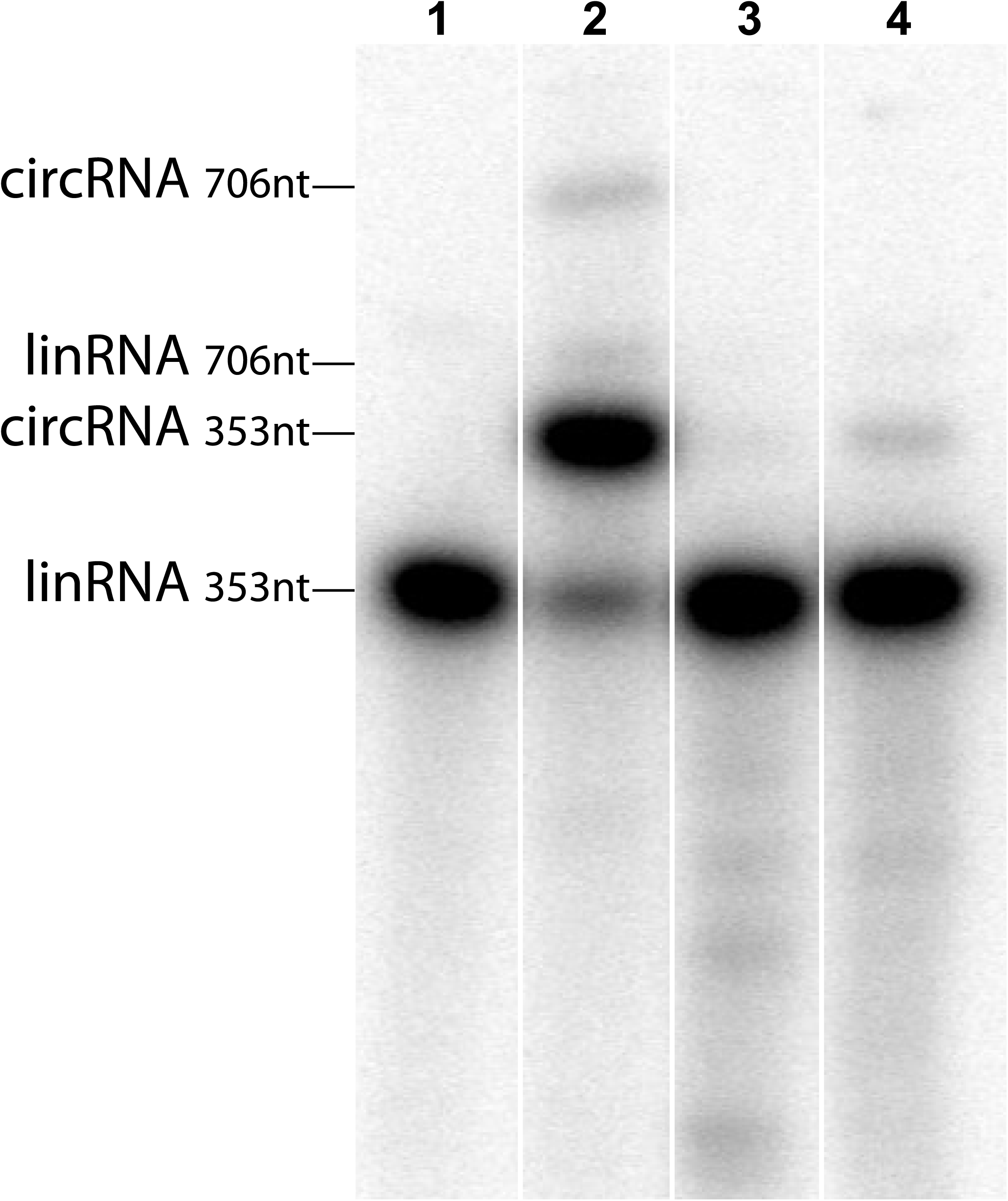
Circularization of Mexican axolotl self-cleaved retrozyme RNA. Purified linear monomeric RNA resulting from double self-cleavage of DRtz353 (see Figure 4C) was run directly in a denaturing gel (lane 1), after 1h incubation with RtcB ligase in its corresponding buffer (lane 2), in RtcB buffer without protein ligase (lane 3), and in 50 mM Mg^2+^ buffer (lane 4). Circular RNAs can be readily detected in lane 2 (up to 80% circularization for the 350 nt monomer) and lane 4 (3% self-circularization for the 350 nt monomer). Minor fractions of linear and circular dimeric RNAs are also indicated (lane 2).

## DISCUSSION

The results reported in this work answer a longstanding question about the presence and role of the paradigmatic HHR in diverse animal genomes. The first HHR motif described in a metazoan genome was reported more than 30 years ago in the satellite DNA repeats of newts (14). Although further characterization of these tandem repeats in newts and salamanders indicated that they were expressed as monomeric and multimeric self-processed linear transcripts (39), we have now revealed that a larger fraction of the RNA population is in fact composed of circular molecules. Moreover, we also show that repeats with HHRs in the genomes of invertebrates from different phyla are also expressed and efficiently self-processed to result in monomeric linear and circular RNAs. Altogether, these data allow us to conclude that short tandem DNA repeats with HHRs in metazoans, plants (26) and, likely, eukaryotes in general, are a peculiar group of mobile genetic elements that are expressed as linear and circular RNAs, probably in most cell types. Moreover, the differences observed in the ratio of linear to circular molecules among diverse species or RNA extracts (see above), allow us to suggest that the levels of circRNAs *in vivo* could be much higher than their linear counterparts. The observed levels of linear RNAs may result from either HHR self-cleavage or spontaneous breakage of the circRNAs during their purification and manipulation. Moreover, with the exception of the predicted secondary structure of the mussel circRNA retrozyme, RNA circles of most metazoan retrozymes are predicted to allow the HHR to freely adopt the catalytically competent structure of the ribozyme (Figure 2C, 2D, 4E and supplementary Figure S6). This feature, observed for most retrozyme circRNAs of metazoans, contrasts with what is observed in plant LTR-retrozymes and viroidal circRNAs with HHRs, which show the ribozyme motif paired with a complementary sequence that should avoid its self-cleavage (26). Consequently, most circRNAs derived from metazoan retrozymes may follow *in vivo* different mechanisms to control their HHR activity and circRNA self-cleavage.

As previously hypothesized (26), plant and animal retrozymes would follow a similar life cycle for their spreading throughout genomes as non-autonomous retroelements. Both types of retrozymes express and accumulate at high levels in either somatic or germinal tissues of any organism analysed, an atypical behaviour for retroelements, which are generally inactive in most cell types and conditions (44-46). But eukaryotic retrozyme repeats are intensively transcribed to produce oligomeric RNAs, which self-process through their HHR motifs. The resulting monomeric RNAs carrying 5’-hydroxyl and 2’-3’-cyclic phosphate ends may undergo covalent circularization by host RNA ligases. These abundant retrozyme circRNAs in the cell can be regarded as stable retrotransposition intermediates and, among other uncertain roles for such naked RNAs or ribonucleoprotein complexes (47), they would be the templates for RTs encoded by autonomous retroelements. This mechanism requires that a properly primed circRNA (either DNA target-primed or through a small cellular RNA) should be recognized by the RT to produce tandem cDNA copies that would be integrated in new genomic loci (supplementary Figure S9). In plants, LTR retrozymes are likely mobilized by the machinery of LTR retrotransposons of the Gypsy family (26). On the other hand, metazoan retrozymes, which lack any of the typical LTR features (Figure 1B and C), should be mobilized by some other active member of the families of non-LTR retrotransposons, such as LINEs or PLEs among others. Interestingly, metazoan retrozymes and PLEs share some peculiarities, such as the presence of type I HHRs (23), their occurrence as tandem copies (48), and their co-existence in all the metazoans analysed, which suggest that autonomous PLEs are a strong candidate to complete retrozyme mobilization.

In summary, the presence of these minimal retroelements in metazoans confirms the existence of a new natural pathway for circRNA biosynthesis in animals through autocatalytic RNAs, in addition to the already described circRNA biogenesis through backsplicing (49). Future research on RNA circles with self-cleaving ribozymes will help us to decipher the roles of these abundant molecules among animals, plants and eukaryotes in general, but also to develop new biotechnological tools in the emerging field of circRNAs.

## DATA AVAILABILITY

All data are available from the corresponding author upon request. The circRNA sequences cloned from *M. galloprovincialis* have been deposited in the GenBank database under the accession codes MN642551-MN642576.

## SUPPLEMENTARY DATA

Supplementary Data are available at NAR Online.

## ACKNOWLEDGMENT

We acknowledge the excellent technical assistance of M. Pedrote.

