## Supplemental Figures S1 to S9 and legends for "Small circRNAs with self-cleaving ribozymes are frequently expressed in metazoan transcriptomes"

**Legends for Supplementary Figures.**

**Supplementary Figure S1**

A selected group of the major type I HHR families detected in the genome of the coral *A. millepora*. Family 1 motifs of (boxed inset) correspond to the canonical type I HHR present in retrozymes (~200 bp repeats), whereas all the other families of minimal HHRs are related to autonomous retrotransposons, such as PLEs or Terminons. The consensus sequence (70% homology) of the aligned HHRs for each family is shown. Only those sequences allowing a perfect helix topology and showing the conserved catalytic center (CUGANGA / GAAAY / RUH) were used for the HHR alignments. Nucleotides in red correspond to conserved non-canonical (47) loop-loop interactions of type I HHRs.

**Supplementary Figure S2**

(A) Sequence alignment of selected retrozyme elements from 12 different species of stony corals. The first position of each sequence corresponds to the self-cleavage site of the retrozyme HHRs. Highest sequence homology occurs among either the ribozyme motifs or the central region of the retrozymes (positions ~100-150 nt). (B) Minimum free energy secondary structure predictions of circRNAs derived from genomic retrozymes sequences from diverse stony corals. These elements were found either in this work or hypothesized in our previous publication (e. g. *A. digitifera* and *P. strigosa*) (26). Sequences corresponding to the hammerhead ribozyme are shown in purple.

**Supplementary Figure S3**

Minimum free energy secondary structure of a retrozyme circRNA sequence predicted from the genome of the anemone *Nematostella vectensis*. The sequence corresponding to the hammerhead ribozyme is shown in purple.

**Supplementary Figure S4**

Consensus sequences of two type I HHRs detected in the retrozymes of the *Mytilus galloprovincialis* genome (HHR A and HHR B consensus results from 2,880 and 1,681 motifs respectively). Both motifs drawn on the left correspond to essentially the same sequence, except for changes in loops 1 and 2 (nucleotides in red), and the upper stem of the helix I (nucleotides in grey). (R: A or G, Y: C or U, W: A or U, K: G or U)

**Supplementary Figure S5**

Sequence alignment of cloned circRNAs from the gonads of two male mussels (*M. galloprovincialis*). The first position of each sequence correspond to the HHR self-cleavage site.

**Supplementary Figure S6**

Minimum free energy secondary structure of some circRNAs encoded by putative retrozymes in the genomes of invertebrates from different phyla, such as (A) rotifers, (B) trematodes, (C) insects, (D) annelids, (E) planarians or (F) crustaceans.

**Supplementary Figure S7**

(A) Sequence alignment of two pairs of retrozyme sequences from the axolotl (*Ambystoma mexicanum*) genome. Each pair corresponds to the Rtzm330 (G327 and G331) and Rtzm350 (U348 and U351) families. (B) Consensus sequences of four representative examples of minimal HHRs characteristic of PLE and Terminon retrotransposons detected in the axolotl genome.

**Supplementary Figure S8**

(A) Autoradiography of self-ligation experiments performed with a doubly self-cleaved linear monomeric retrozyme from *M. galloprovincialis* obtained from *in vitro* transcription of a dimeric construct (DRtzm375_Mg). Experiments were performed in the absence of $Mg^{2+}$ (lane 1), with 10 mM $Mg^{2+}$ (lane 2) and with 50 mM $Mg^{2+}$ (lane 3). Dimeric, circular and linear RNA molecules are indicated. (B) Ethidium bromide-stained gel showing circularization experiments of the retrozyme monomer Rtzm375_Mg in the absence (lane 1) and in the presence (lane 2) of RtcB ligase. A detail of the lane 2 of the gel after silver staining is shown on the right.

**Supplementary Figure S9**

A tentative model for the replication cycle of metazoan (non-LTR) and plant (LTR) retrozymes. Either non-LTR retrozymes carrying type I HHRs described in metazoans or LTR retrozymes carrying type III HHRs described in plants could follow a similar propagation pathway through circRNAs as retrotransposition intermediates, which would be reverse transcribed as oligomeric cDNA repeats, and integrated at a new genomic locus by the machinery of autonomous retrotransposons present in the host genome.

### Type I HHRs in the genome of the coral *Acropora millepora*

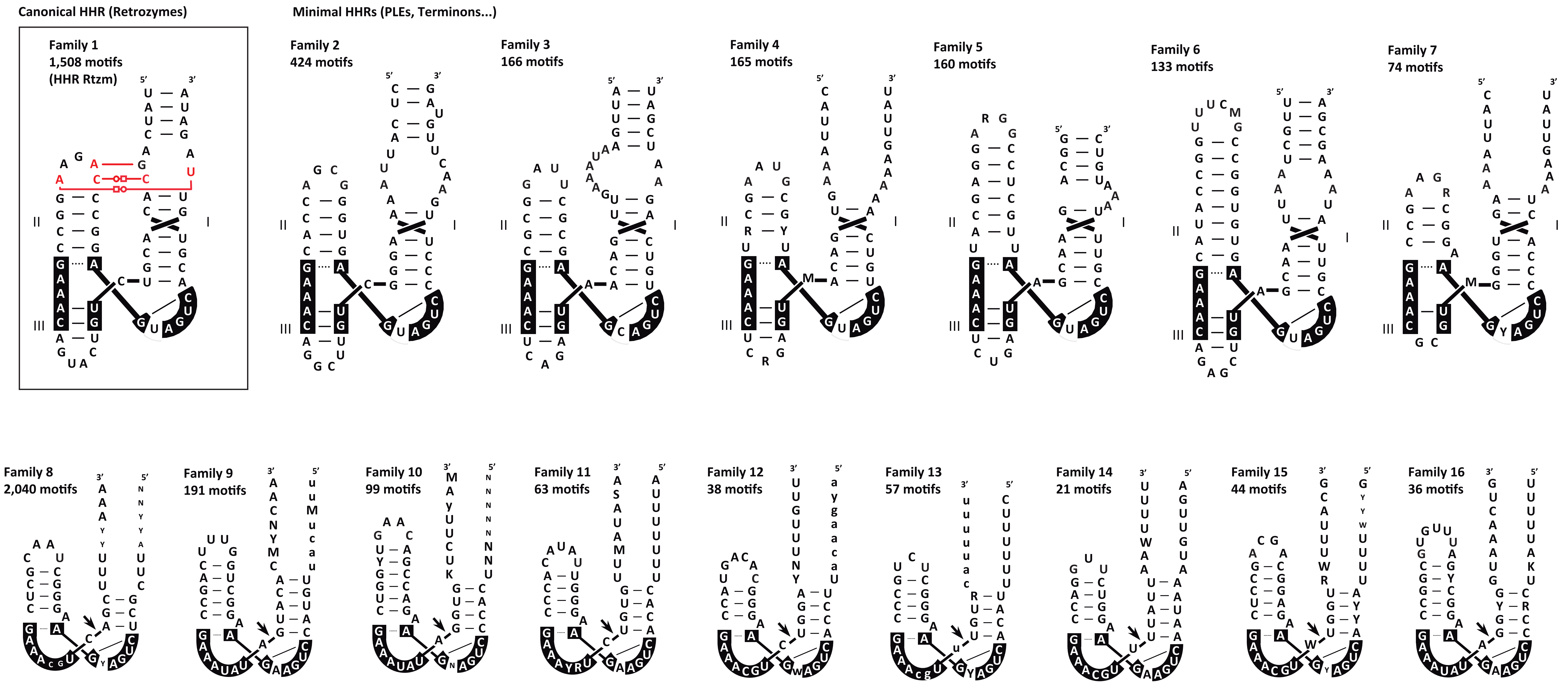


**Supplementary Figure S1**

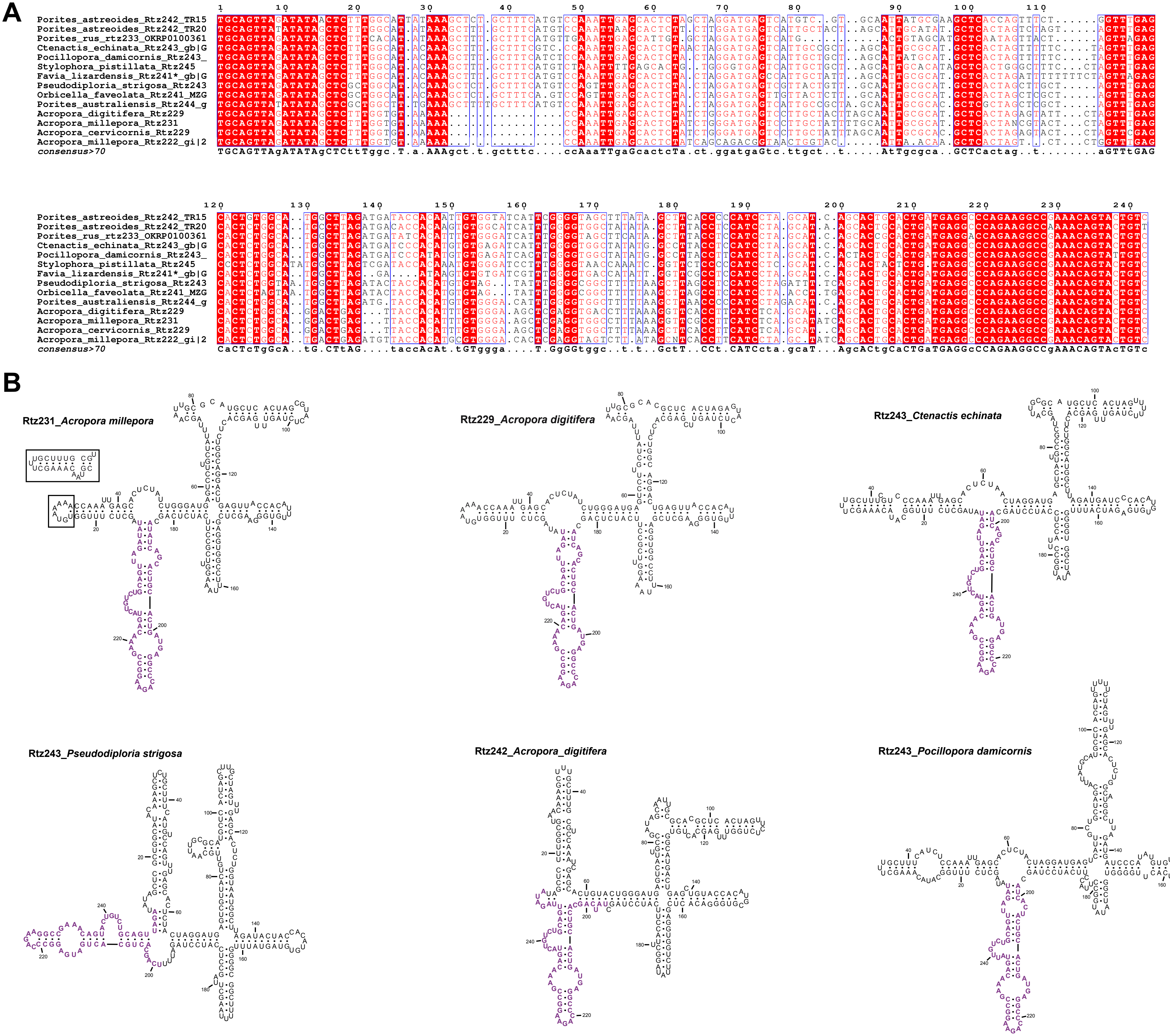


### Supplementary Figure S2

**Rtz354_*Nematostella vectensis***

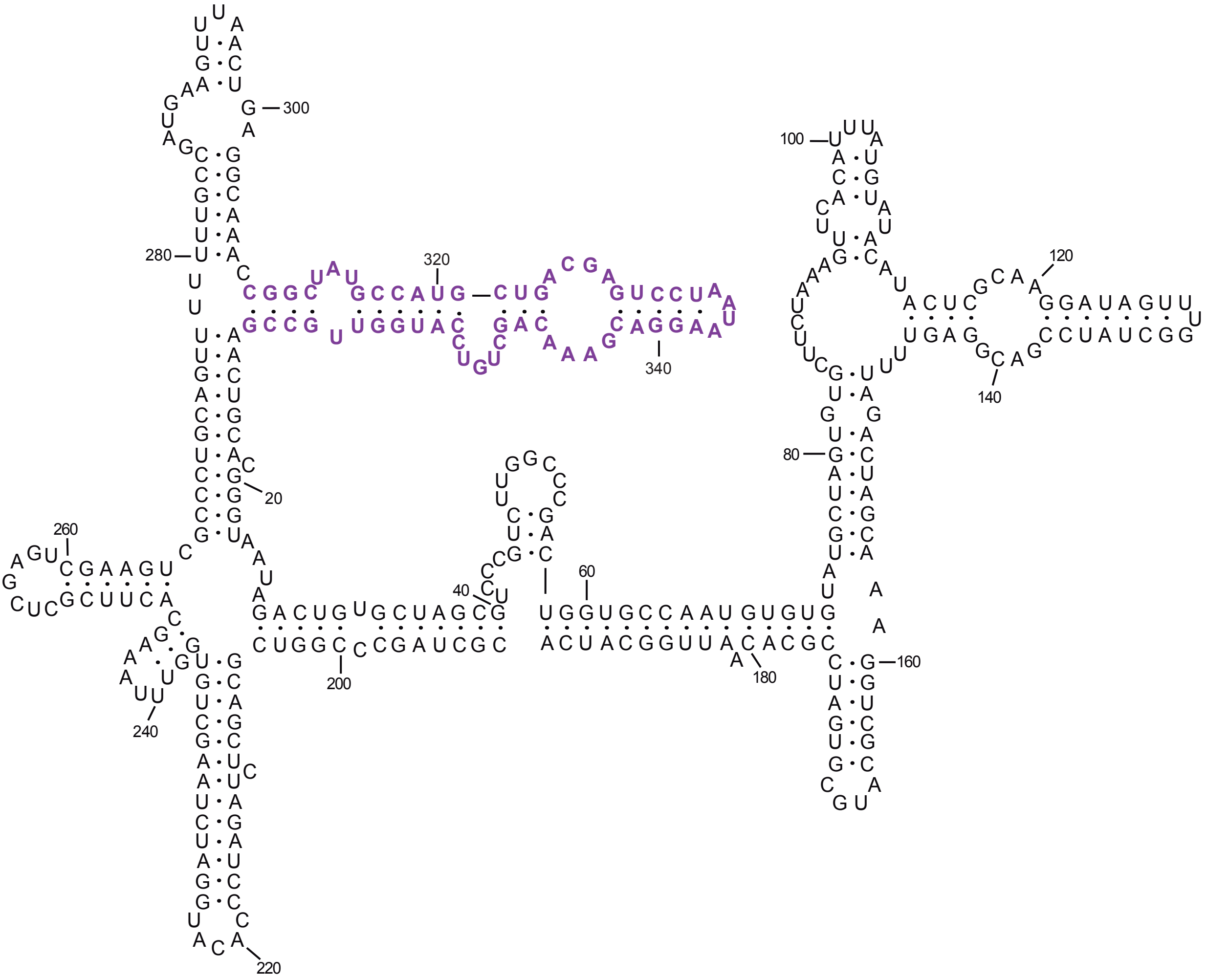


#### Supplementary Figure S3

**HHR A**

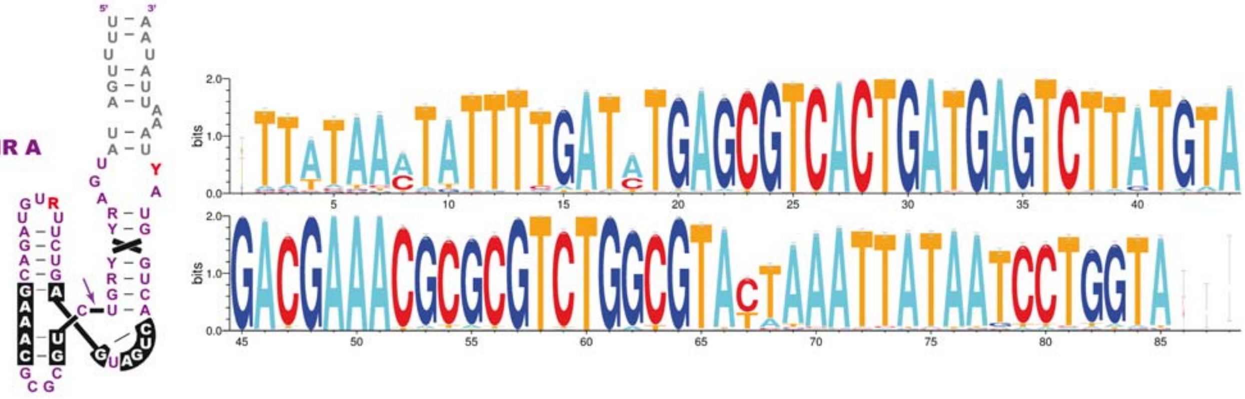


**HHR B**

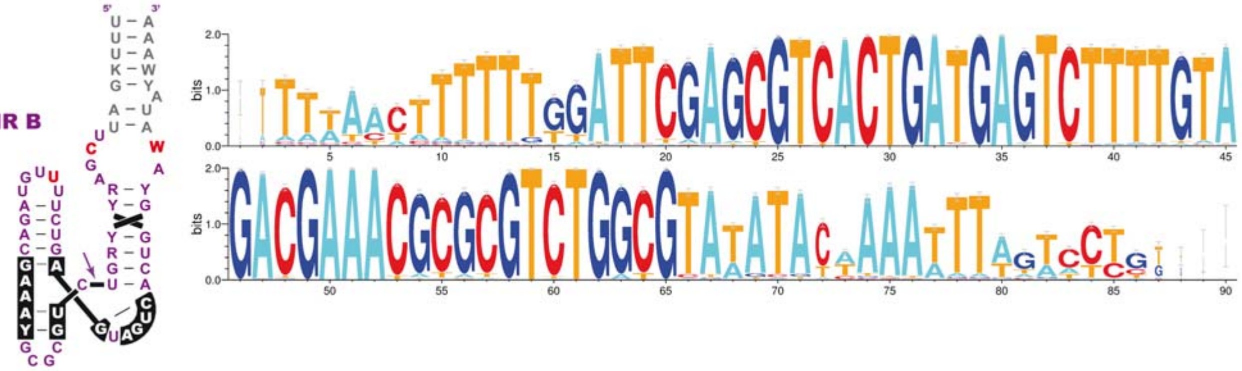

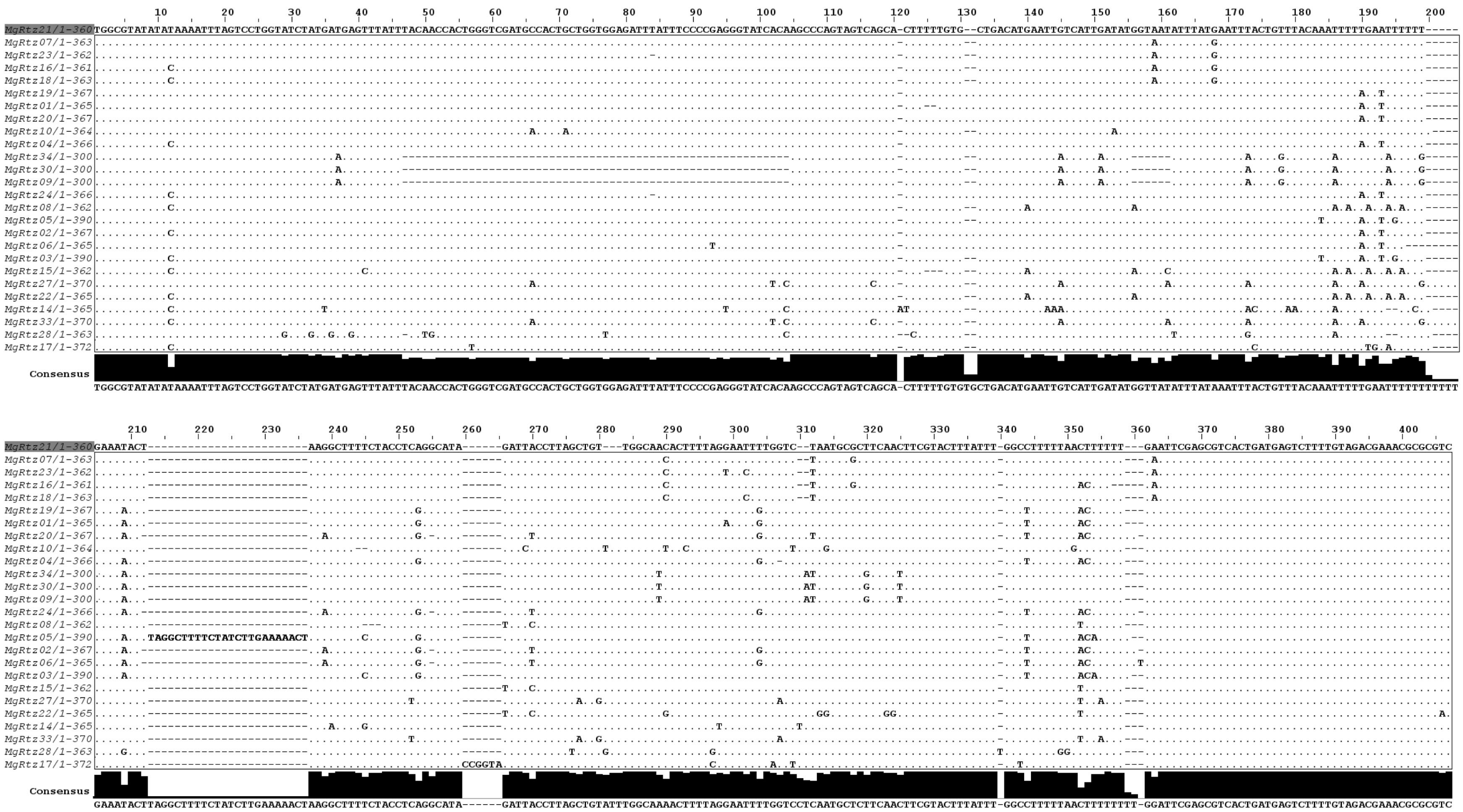


**Supplementary Figure S5**

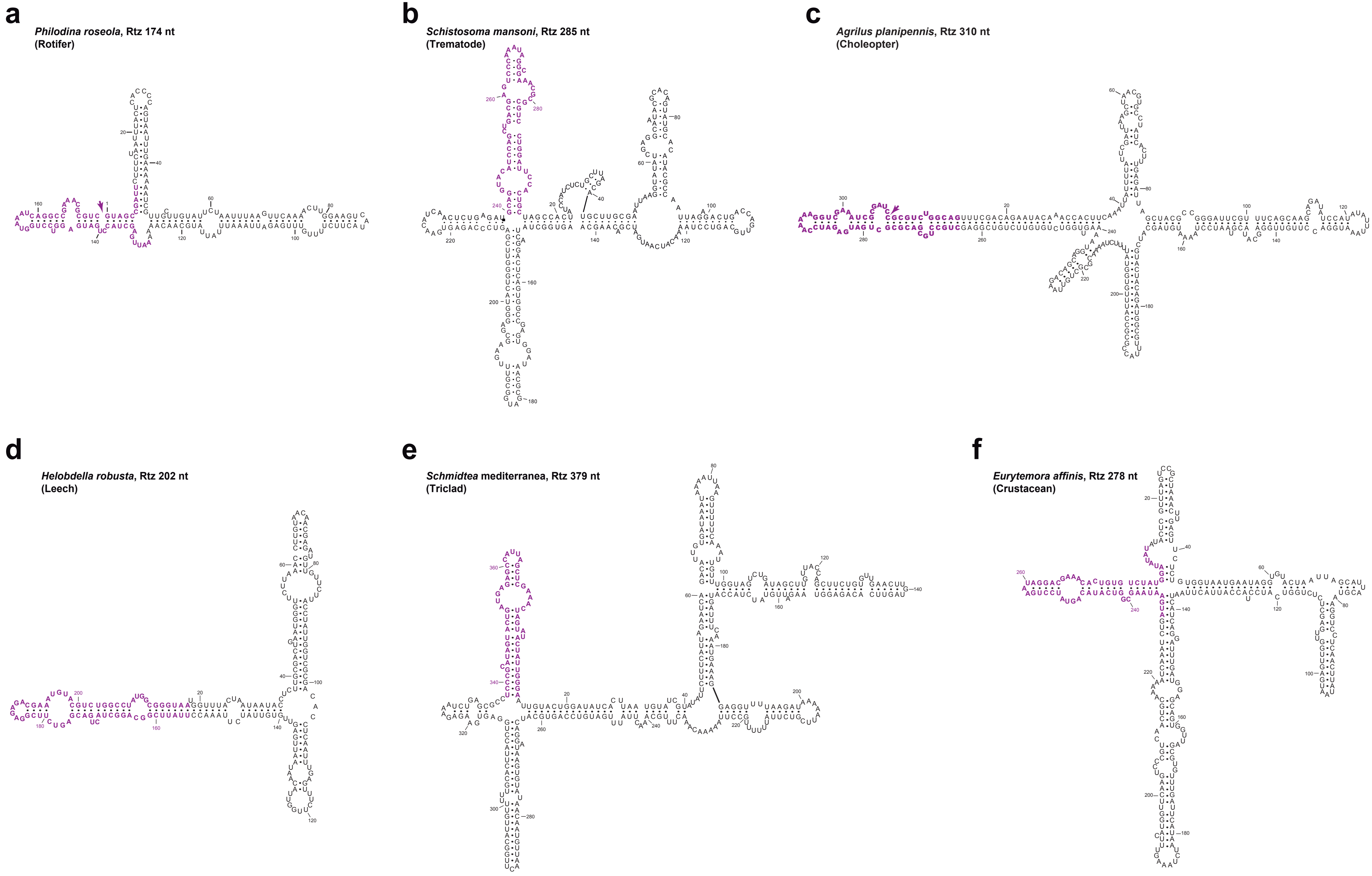


**Supplementary Figure S6**

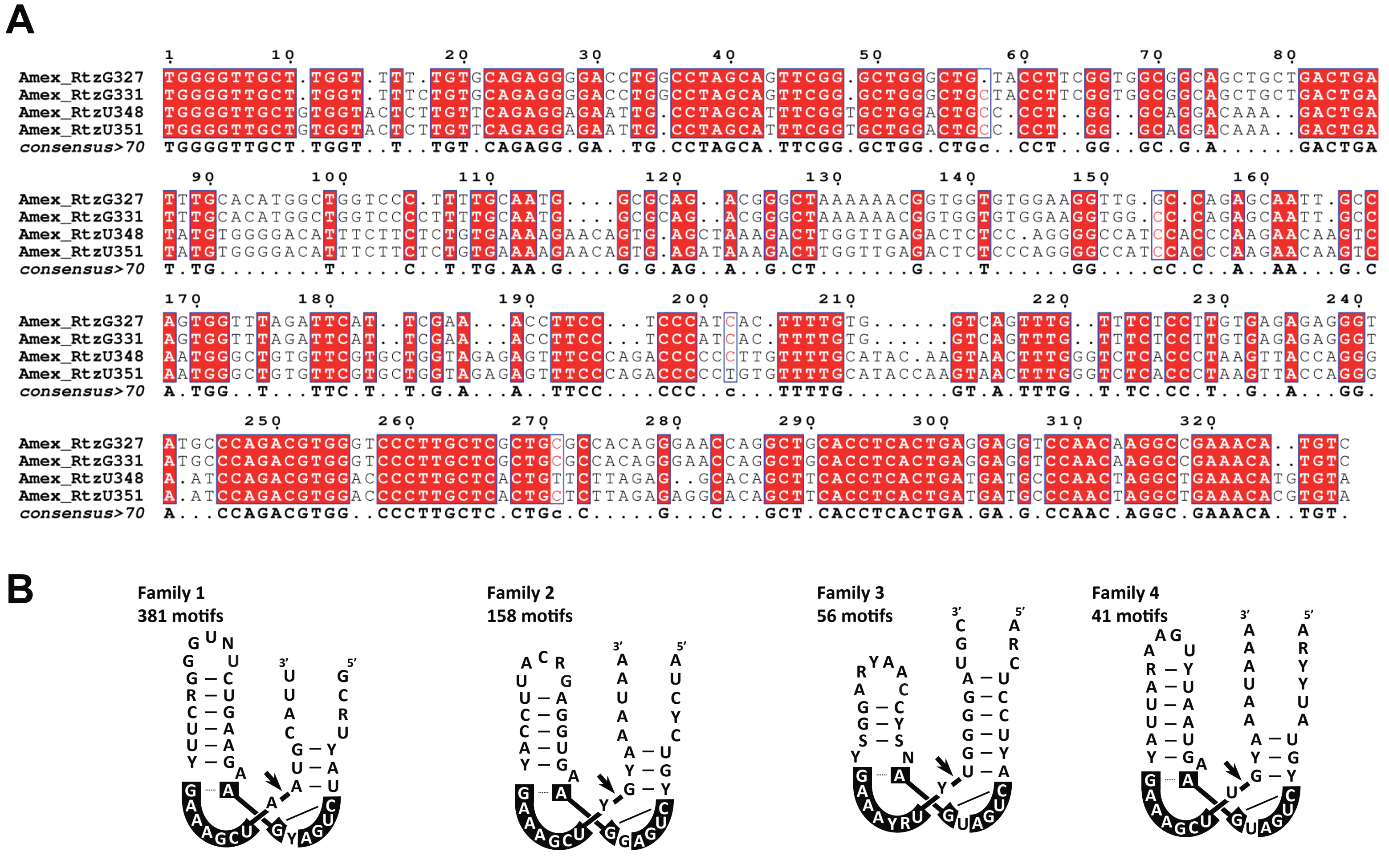


**Supplementary Figure S7**

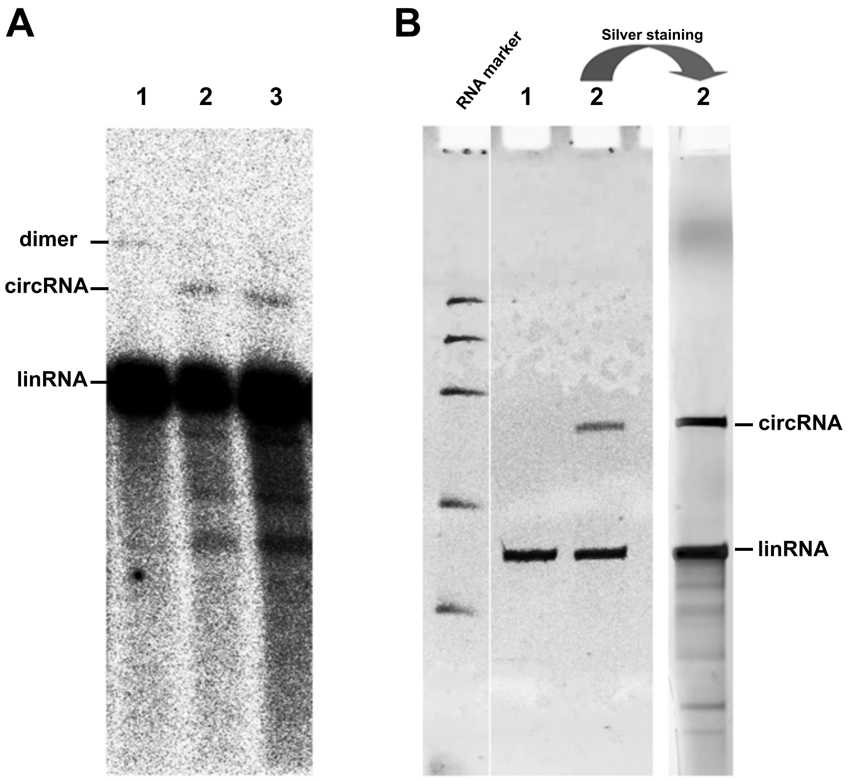


Supplementary Figure S8

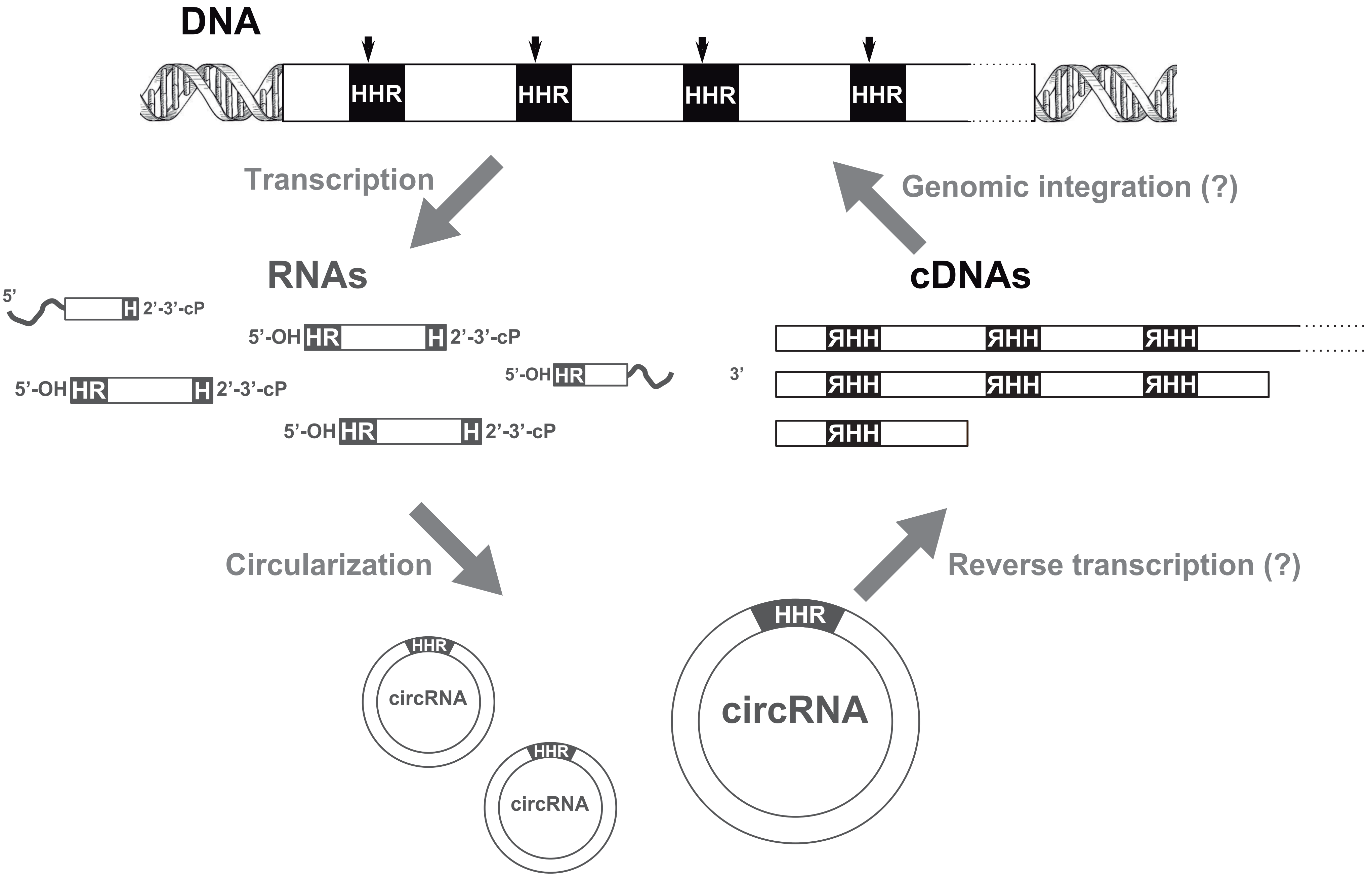


**Supplementary Figure S9**
